# The ICP22 protein of Herpes Simplex Virus 1 promotes RNA Polymerase II activity on Viral Immediate Early Genes

**DOI:** 10.1101/2021.05.05.442792

**Authors:** Claire H. Birkenheuer, Joel D. Baines

**Author notes:** Address correspondence to Joel D. Baines.

## Abstract

To determine the role of herpes simplex virus (HSV-1) ICP22 in viral transcription we performed precise nuclear run-on followed by deep sequencing (PRO-Seq) to map active RNA polymerase II (Pol II) on viral and cellular genomes in cells infected with a viral mutant lacking the entire ICP22-encoding α22 (US1/US1.5) gene, or a virus derived from the deletion mutant but bearing a restored α22 gene. At 3 hours post infection (hpi), the lack of ICP22 reduced Pol II activity at promoter proximal pause (PPP) sites on the α4 and α0 genes, and on the bodies of the α4, α0, and α27 genes. The decreased activity at α0 and α4 PPP sites at 3 hpi was distinguishable from effects caused by treatment with a viral DNA polymerase inhibitor. The ICP22 mutant had multiple defects at 6 hpi, including lower viral DNA replication and reduced Pol II activity on viral genes of all temporal classes. Between 3 and 6 hpi the repair virus, like wild type HSV-1, redirected Pol II activity from cellular genes to viral genes. In the absence of ICP22, the opposite occurred inasmuch as Pol II activity returned from the viral genome to cellular genes. These data indicate that ICP22 acts to increase Pol II activity at the PPP sites and bodies of viral immediate early genes at early times post infection, and directly or indirectly helps retain Pol II activity on the viral genome later in infection.

**Importance:** Using a mutant lacking the full US1/US1.5 gene, this study establishes a role for ICP22 in increasing Pol II activity on viral immediate early genes by enhancing transcription initiation and/or elongation onto immediate early gene bodies. Unlike truncation mutants studied previously, the full null virus is unable to sustain DNA replication and Pol II activity on late genes, precluding later stages of the viral life cycle.

## Introduction

HSV-1 expresses its genes in a coordinated temporal pattern. Initiation of viral transcription occurs when the *α* transinducing factor (*α*-Tif or VP16) enters the cell with the incoming viral particle. The *α*-Tif binds to immediate early, or *α*-gene, promoters and recruits transcription factors and RNA Polymerase II (Pol II) to stimulate expression (1, 2). The *α* proteins then promote expression of *β* genes (2), which encode proteins required for viral DNA replication (2, 3). DNA replication augments the expression of the leaky late *γ*_1_ genes and is necessary for expression of true late or (*γ*_2_) genes encoding mostly structural proteins (4-7).

Pol II transcribes cellular genes in a manner that is mostly conserved from yeast to higher eukaryotes (8). First, Pol II is recruited to the gene promoter by transcription factors bound to the gene (8). Once assembled at the promoter, cyclin dependent kinase 7 (CDK7) phosphorylates the carboxy terminal domain (CTD) of Pol II at serine 5 in its heptad repeat (9) to initiate transcription at the transcription start site (TSS). On many host genes, transcription continues for approximately 50 nucleotides producing a short nascent RNA before undergoing promoter-proximal pausing (PPP). Pol II remains paused until serine 2 in its CTD heptad repeat region is phosphorylated by CDK9 (9, 10). This phosphorylation event is associated with release of Pol II from the PPP site followed by transcription through the downstream gene body until encountering a transcription termination signal (TTS) (9).

An emerging picture suggests that HSV-1 transcription mimics certain aspects of this model for its own transcription. For example, chromatin immunoprecipitation (ChIP) experiments show that Pol II accumulates around the 5’ end of viral genes and pauses downstream of TSS on many viral genes (7, 11). Inconsistent with this model, however, is the observation that Pol II phosphorylation at serine 2 of the CTD is reduced in infected cells and ICP22, a ∼68-kDa ICP22 immediate early protein encoded by the *α*22 or US1 gene, is necessary and sufficient to mediate this change (12-17). This ICP22 activity is especially puzzling because (i) ICP22 has been implicated in enhancing, not inhibiting, transcriptional elongation on viral genes, and (ii) CDK9 activity is essential for viral replication despite the fact that ICP22 reduces the Pol II Serine 2 phosphorylation that CDK9 mediates (11, 12, 15, 16, 18-21). ICP22’s role in regulating transcription elongation is supported by its ability to recruit the FACT complex, also known as the host chromatin transcription complex, which is composed of SSRP1 and SPT16, and its associated factor SPT6 to viral genomes (11). Moreover, ChIP experiments performed with different antibodies towards Pol II have shown that deletion of ICP22 at 6 hpi causes a reduction of Pol II in viral gene bodies, but similar levels of Pol II at the 5’ end of the genes suggesting that ICP22 plays a role in regulating Pol II transcription elongation, but not Pol II initiation (11).

PRO-seq is a deep sequencing technique that takes advantage of the inhibitory nature of biotinylated nucleotide incorporation into nascent RNA during a nuclear run-on reaction (10, 22). Once incorporated into the nascent RNA, biotin replaces the free 3’ hydroxyl to prevent Pol II from further transcript extension. By performing the nuclear run-on in the presence of a high concentration of biotinylated nucleotides, Pol II’s location can be deduced by sequencing the nascent biotinylated RNA and mapping its 3’ end to the template (22).

Using PRO-seq we previously demonsthrated that Pol II pauses upstream of many viral gene bodies at approximately 50 nt downstream of the TSS (7). Furthermore, the level of active Pol II is decreased on the viral genome when viral replication is repressed with either the CDK9 inhibitor flavopiridol, or viral DNA replication inhibitors acyclovir and phosphonoacetic acid (PAA) (7). Moreover, blocking viral DNA replication significantly reduces signal in standard nuclear run-on reactions (15), and leaves only small amplitude PPP peaks on viral genes at 6hpi (7). Taken together, these observations suggest that release from PPP and full occupancy of active Pol II on the viral genome are regulated by CDK9 and viral DNA replication (7, 13, 14, 18, 23).

In this study we used PRO-Seq to map the location of transcribing Pol II on the viral and host genome at 3 and 6-hours post infection (hpi) in the presence and absence of ICP22. The resulting data indicate an important role for ICP22 in ensuring optimal Pol II activity on immediate early genes early in infection, resulting in substantial defects in replication later in infection.

## Results

### PRO seq analysis of Viral genes

To determine the role of ICP22 in viral transcription we infected cells with a previously described virus lacking the entire *α*22 (US1/US1.5) open reading frame and a virus derived from that deletion virus but bearing a genetically restored α22 (21, 24). Thus, phenotypic differences between the two viruses should be ascribable to the absence of *α*22 and its encoded ICP22 protein. The infected HEp-2 cells were subjected to PRO-seq analysis at 3 and 6 hpi as previously described (7, 25). Read counts were normalized to spiked in drosophila DNA to ensure consistency between biological replicates.

We first looked at Pol II distribution across the entire viral genome using the Integrative Genomics Viewer (IGV) (26) from the Broad Institute (Fig 1A). Consistent with our previous observations using wild type virus, overall Pol II activity on the genome of the genetically repaired virus was high at 3 hours and increased from 3 to 6 hpi (blue and gold data, respectively) (Fig 1A). In contrast, Pol II activity did not appear to increase on the ΔICP22 genome over this time period (green and magenta) (Fig 1A). When we quantified all normalized reads aligning to the viral genome from each infection using SeqMonk software (27) we found no statistical difference between the total amount of active Pol II on the two viral genomes at 3 hpi. However, significantly less active Pol II was associated with the ΔICP22 viral genome than with the repaired viral genome at 6 hpi (Fig 1B). These data indicate that ICP22 is required to increase occupancy of active Pol II on the viral genome between 3 and 6 hpi.

**Figure 1.**
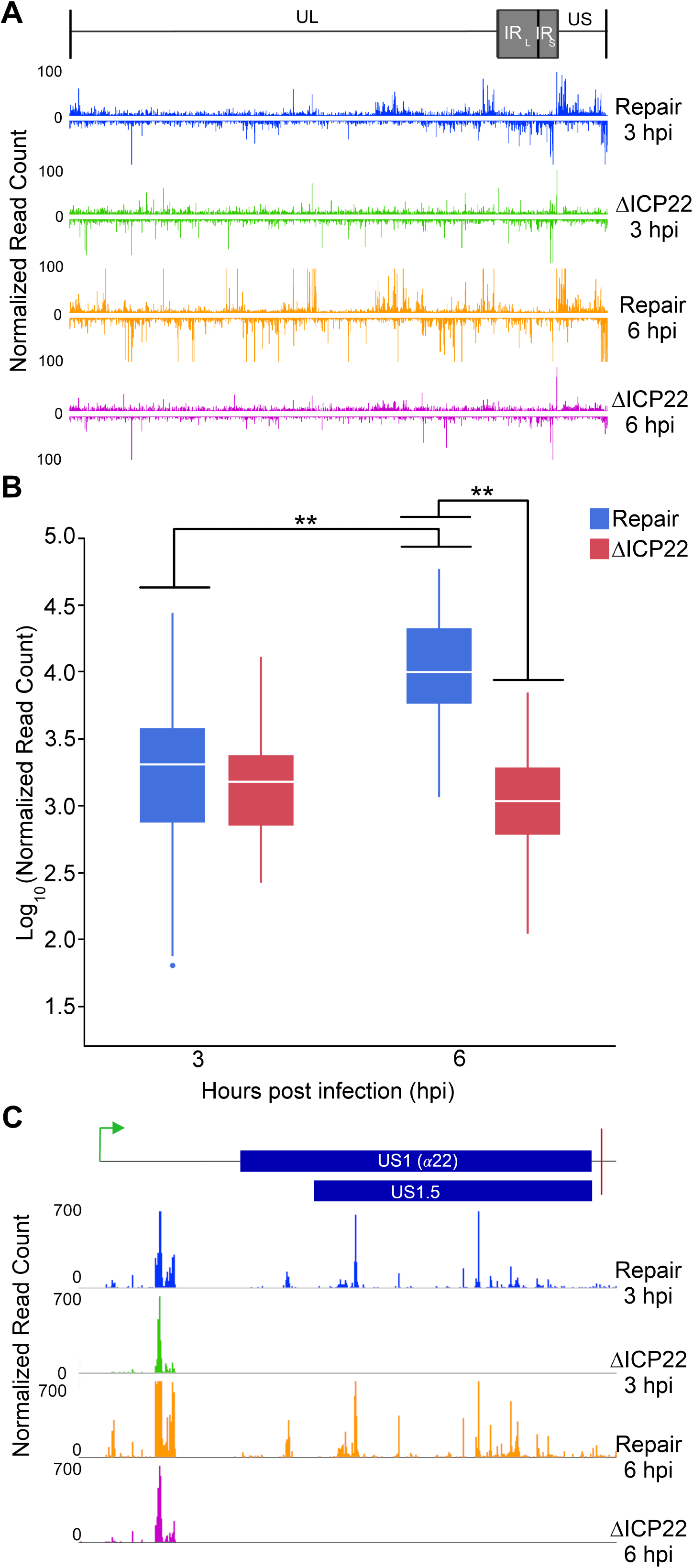
(**A**) The normalized read counts from a representative PRO-seq replicate aligned to the HSV-1 (F) genome. The diagram at the top shows the location of the unique long (UL), unique short (US), the long internal repeat (IR_L_), and the short internal repeat (IR_S_) in the viral genome. External repeat sequences have been removed. The ΔICP22 data is shown in green and magenta at 3 and 6 hpi, respectively. Repair virus data are shown in blue and gold at 3 and 6 hpi, respectively. Reads matching both strands of the double stranded HSV-1 genome are shown, with genes running left to right on the top and right to left on the bottom. One normalized read count is the equivalent to one actively transcribing Pol II molecule. (**B**) The log_10_ transformed normalized read count values from the entire HSV-1 genome from all replicates at 3 and 6 hpi in both the repair and ΔICP22 infections. The data is shown in a box and whisker plot with the mean indicated by the white line in the middle of the box. Statistical analysis was done using a two-way ANOVA with a Tukey’s honestly significant difference (HSD) post-test. A ** indicates a p-value ≤ 0.0001. (**C**) Normalized PRO-seq reads from each indicated experimental set matching the *α*22 (US1/US1.5) gene. Only the *α*22 coding strand (upper) is shown. The collinear locations of the US1 and US1.5 open reading frames (blue rectangles), the *α*22 transcription start site (green arrow), and *α*22 transcription termination signal (red line) are shown.

Using IGV and the normalized read count values, we verified that the *α*22/US1/US1.5 orfs encoding ICP22 was deleted in the ΔICP22 virus inasmuch as it produced no PRO-seq reads from the *α*22 open reading frame during the time course examined (Fig 1C) (21).

To analyze the level of Pol II occupancy on viral genes of different temporal classes, we compared Pol II activity on genes of both viruses (TSS to TTS) that have a well-accepted temporal class designation (7). Average levels of Pol II activity on the genes of the repair virus (black bars) and ΔICP22 virus (gray bars) are shown in Fig 2 A-D. As expected, the Pol II levels on the immediate early *α* genes of the repair genome stayed consistent and high from 3 to 6 hpi (Fig 2A). While Pol II activity on the later gene classes of this virus were lower than were present on the *α* genes at 3 hours, these levels increased at 6 hours (Fig 2 B-D). These observations indicate that Pol II occupancy on repair virus genomes increases over time similar to that of the wt virus as previously described (7).

**Figure 2.**
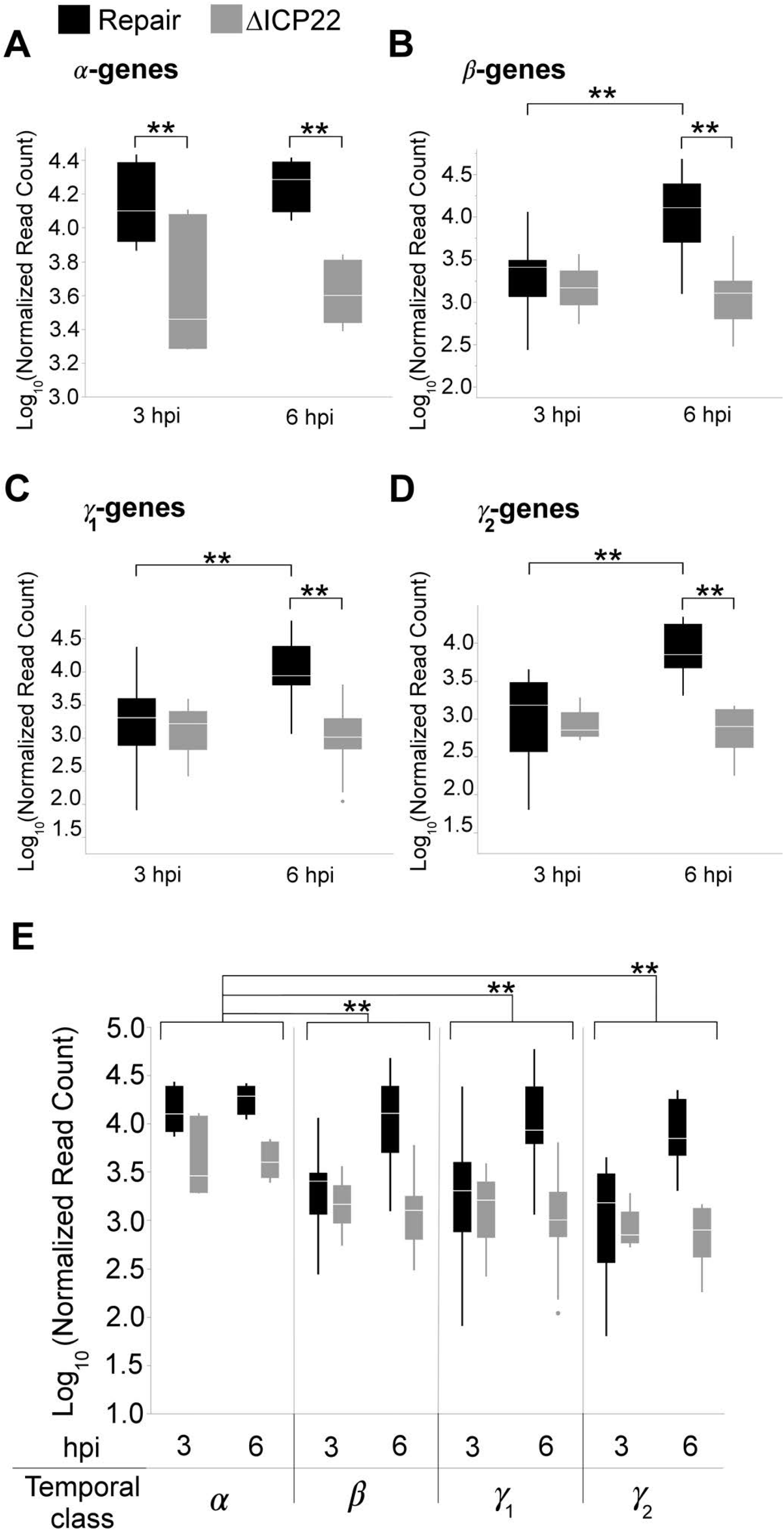
(**A-D**) Normalized PRO-seq data matching different temporal classes of viral genes. Read counts matching the entire gene bodies (TSS to TTS) of individual genes in each kinetic class are shown. Log_10_ transformed read counts for the repair virus infection are indicated in black, and the ΔICP22 values are indicated in gray. A two-way ANOVA was performed comparing the normalized read counts between the two groups over time. Significance was determined using a Tukey’s HSD post-test and a ** indicates a p-value ≤ 0.01. (**E**) A three-way factorial comparison of the data presented in A-D, showing the difference in normalized read counts between the ΔICP22 and repair infections across the temporal classes. A ** indicates a p-value ≤ 0.01 indicating ICP22 deletion affects *α* genes differently than it does genes of other temporal classes.

Importantly, the three analyzed *α* genes in the ΔICP22 virus genome (*α*0, *α*4, and *α*27) bore significantly less Pol II at both 3 hpi, and 6hpi than the same genes of the repair virus (Fig 2A). While Pol II occupancy on viral genes belonging to the *β, γ*_1_, and *γ*_2_ temporal classes (Fig 2 B-D) was similar between the two viruses at 3 hpi, it was lower on these genes in the ΔICP22 viral genome than on the repair viral genome at 6 hpi. These data indicate that ICP22 is responsible for recruiting or maintaining active Pol II onto *α* genes from 0 to 3 hpi, and for increasing active Pol II occupancy on later genes between 3 and 6 hpi (Fig 2E).

### Viral DNA replication in the ΔICP22 virus

Because *α* proteins are required for expression of the viral DNA replication machinery, the significant difference observed in Pol II occupancy on the *β, γ*_1_, and *γ*_2_ gene classes at 6hpi in the ΔICP22 infection (Fig 2E) could be due to decreased *α* gene protein products (Fig 2A), decreased viral DNA replication (7), or a combinatorial effect of the two. To address the role that viral DNA replication plays in the ΔICP22 Pol II phenotype we quantified viral DNA at various times after infection with either the ΔICP22 or repair viruses. As shown in Fig 3, viral DNA replication was inhibited in the ΔICP22 virus infected cells when compared to levels in cells infected with the repair virus beginning at 3 hpi and continuing through the 12-hour time point. The 12-hour samples showed the most significant difference in DNA quantity between the two infections (Fig 3). A defect in viral DNA replication was also noted in studies of Kawaguchi and colleagues in earlier studies of these viruses (21).

**Figure 3.**
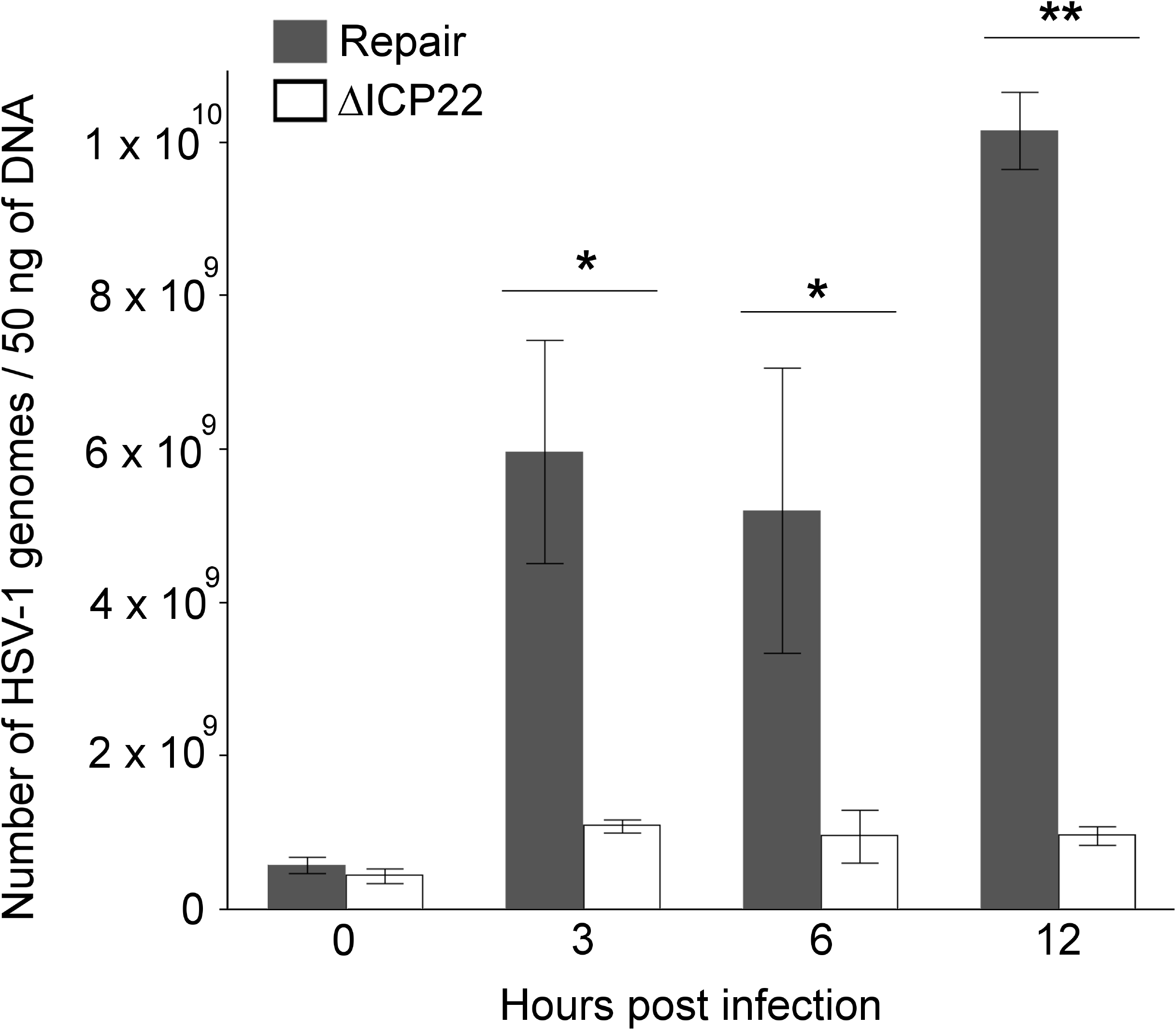
Number of HSV-1 DNA genomes over time in the Repair (gray-filled boxes) and ΔICP22 (white-filled boxes) virus infections. The y-axis indicates the number of HSV-1 genome copies found in 50ng of total DNA extracted from 2 × 10^7^ HEp-2 cells infected at an MOI of 5. The number of HSV-1 DNA genomes was calculated using absolute quantitative PCR, using a standard curve prepared from serial dilutions of a pcDNA3 plasmid containing the entire HSV-1 (F) UL51 sequence. HSV-1 DNA copy numbers were determined for each virus infection from two experimental replicates which are averaged and shown with their corresponding standard deviation in the bar chart. A two-way ANOVA followed by a Tukey’s HSD post-test was performed comparing the DNA levels between the virus infections over time. A * represents a p-value ≤ 0.05, and a ** indicates a p-value ≤ 0.01.

### Pol II occupancy on viral *α* genes in the ΔICP22 virus and repair virus treated with PAA at 3hpi

The above data suggested the possibility that the ΔICP22 PRO-Seq profiles observed at 3 and 6 hours on the ΔICP22 viral genome were due in part to defects in DNA replication rather than a direct effect of the lack of ICP22. To distinguish between these possibilities, we performed PRO-seq on cells infected with either virus in the presence or absence of the viral DNA replication inhibitor phosphonoacetic acid (PAA), and specifically compared the Pol II profile on the three viral *α* genes (TSS to TTS) (Fig 4 A-D). Analysis was not performed on *α*22 because the *α*22 orf was deleted, nor on *α*47 because its 5’ end is located in the external short repeat region of the genome thus precluding clear assignment separate from the *α*22 TSS.

**Figure 4.**
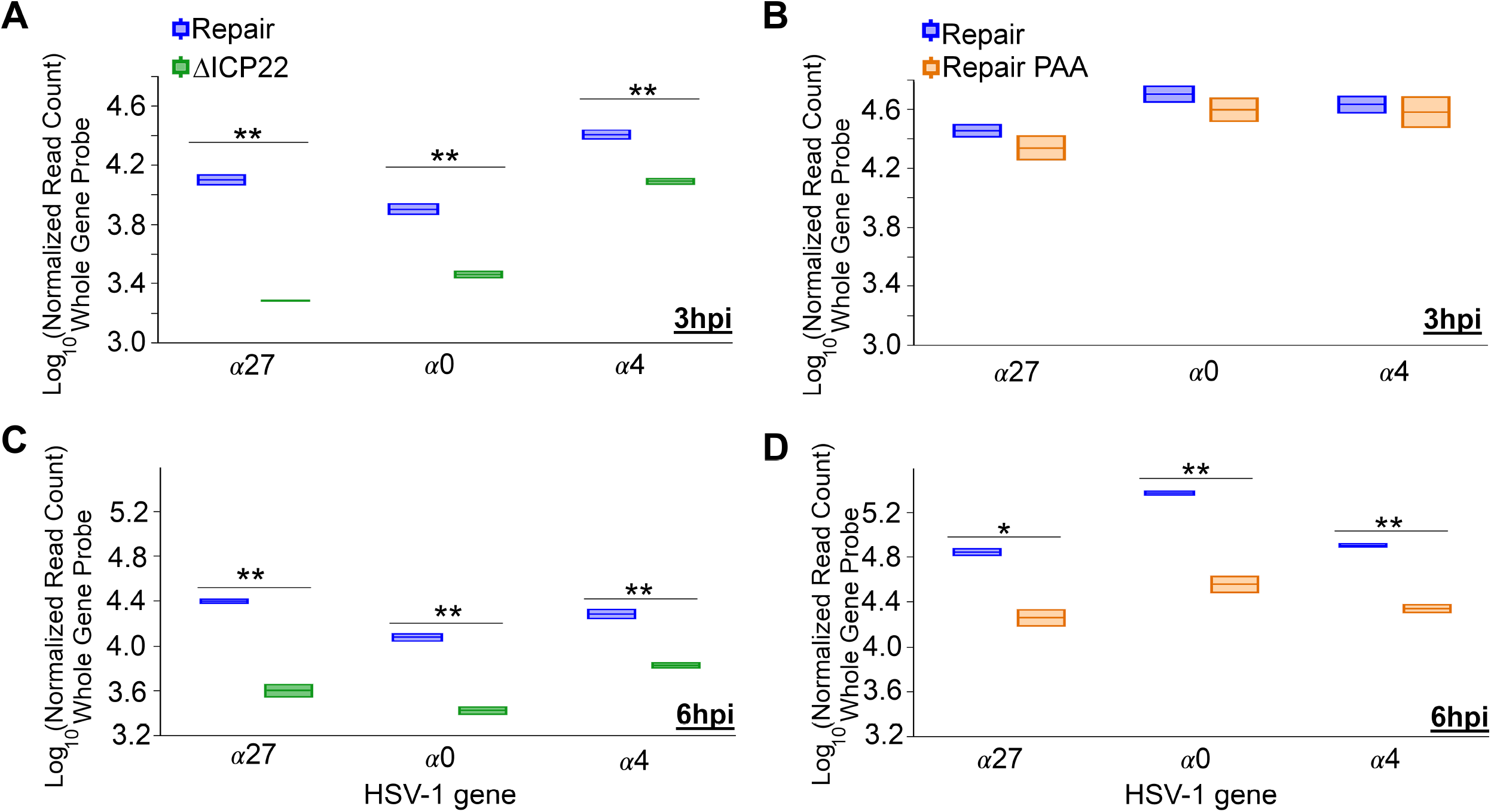
Analysis of normalized PRO-seq read counts on three viral *α* genes. The comparison of the two biological replicates from each of the ΔICP22 (green) / repair (blue) virus infections is shown in **A and C** at 3 hpi (**A**) and at 6 hpi (**C**), and the comparison of the two biological replicates from the PAA treated (orange) and non-treated (blue) repair virus infections is shown in **B and D** at 3 hpi (**B**) and at 6 hpi (**D**). A one-way ANOVA with a pooled T-test was used to compare the Log_10_ transformed normalized values from full length (TSS to the TTS) gene probes for the indicated gene under each condition. A ** represents a p-value ≤ 0.01.

As shown in Figure 4A, all three *α* genes of ΔICP22 (0, 4 and 27) had significantly less active Pol II on their loci at 3hpi than the repair virus. Although DNA was replicating to a limited extent at this time (Fig 3), its inhibition with PAA (7) did not statistically alter the level of Pol II occupancy on these genes below that of the repair virus (Fig 4B). As shown in Figure 4C, at 6 hpi, both the ΔICP22 infection and inhibition of viral DNA replication in the repair virus infection reduced Pol II occupancy on the *α* genes as was observed previously in wt HSV-1 (F) infected cells treated with PAA (7). The observations at 3-hpi indicate that ICP22 regulates *α* gene expression whereas DNA replication did not pay a prominent role at this time. By 6 hours, however, both ICP22 and DNA replication contributed to optimal Pol II occupancy on viral genes.

### ICP22 effect on *α*4 and *α*0 Promoter Proximal Pausing

Since ICP22 regulates Pol II phosphorylation that is associated with PPP release, we sought to compare PPP on immediate early genes in the ΔICP22 and repair virus genomes. We focused on the PPP peaks of *α*4 (Fig 5 A-D) and *α*0 (Fig 5 E-H) originally shown in PRO-seq analyses of the wt HSV-1(F) genome (7). The data were obtained using two independent biological replicates for each sample in each experimental set: the ΔICP22 and repair at 3hpi (Fig 5 A and E) and 6hpi (Fig 5 C and G), and PAA-treated and untreated repair infections at 3hpi (Fig 5 B and F) and 6hpi (Fig 5 D and H).

**Figure 5.**
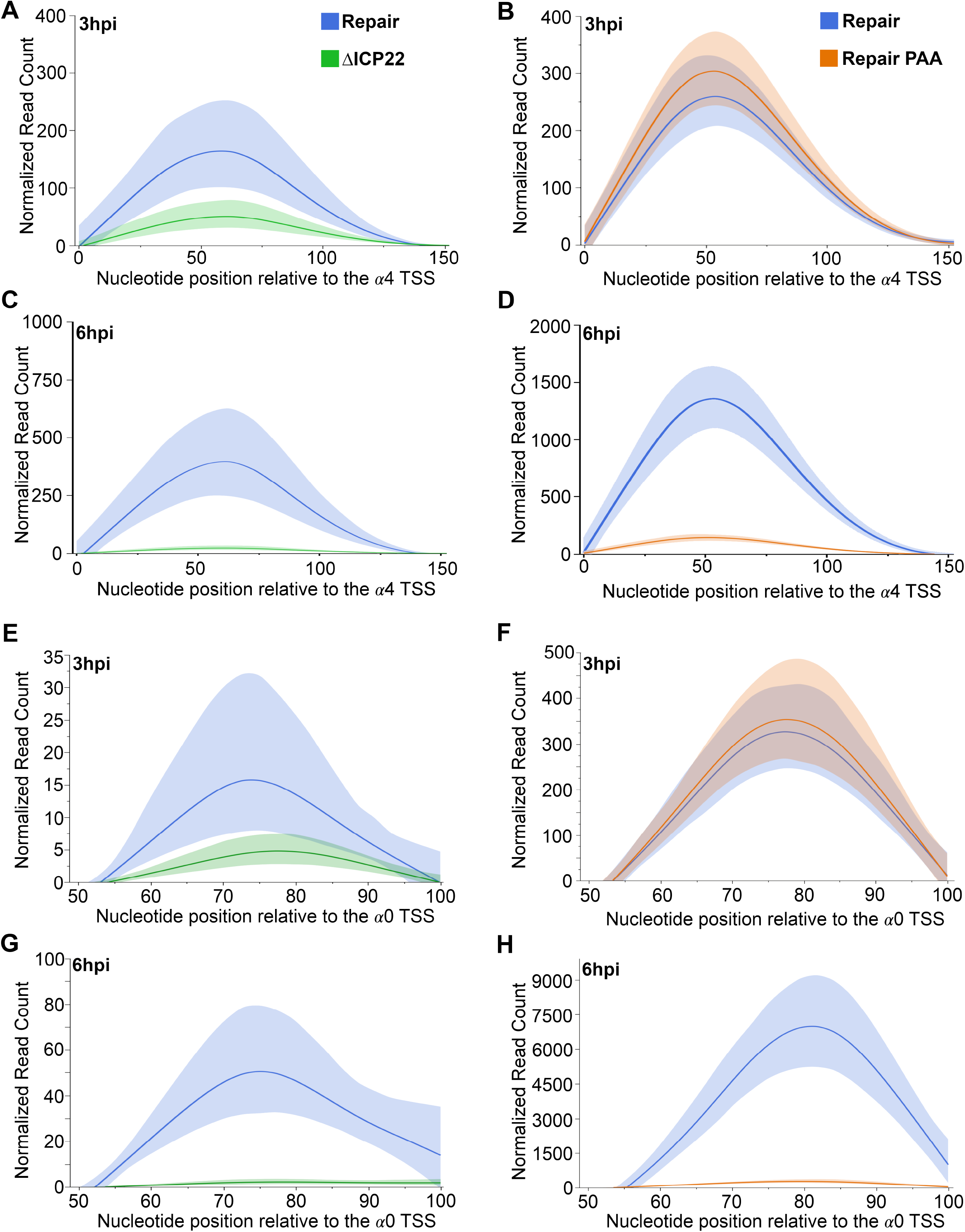
Comparison of the cubic spline interpolations of normalized read counts from each individual nucleotide in the PPP region of the *α*4 gene (**A-D**) and *α*0 gene (**E-H**). Data from infection with the ΔICP22 virus is shown in green, untreated repair virus data is shown in blue, and repair virus treated with PAA is shown in orange. Data taken at 3 hpi (**A, B, E, and F**) and at 6 hpi (**C, D, G H**), are shown. The cubic spline interpolation of the data is shown as the dark line drawn from the data of all experimental sets. The bootstrap confidence of fit for the spline calculation is shown in the lighter, shaded areas and was performed using 300 bootstrap iterations.

The raw read counts from each individual nucleotide spanning the first 200 nucleotides downstream of the *α*4 TSS were normalized to the Drosophila spike-in data, and then used in a cubic spline interpolation to smooth the data to a representative curve shown as a bold line. A bootstrap confidence of fit for the interpolation was calculated using 300 different iterations of the data and is shown as the light-shaded area around the spline (Fig 5).

At 3 hours in the ΔICP22 infection the PPP peaks of *α*4 and *α*0 (green line, Fig 5 A, and E) were reduced in amplitude when compared to the same peaks in the repair virus infection (blue line, Fig 5 A and E). In contrast, the amplitudes of the *α*4 and *α*0 PPP peaks in the repair infection were not affected by PAA treatment (Fig 5 B and F). These observations demonstrate that ICP22 is required to maintain optimal Pol II activity at the *α*4 and α0 PPP sites at 3hpi, whereas blocking DNA replication did not increase or decrease PPP at this time (Fig 3, Fig 4 and Fig 5).

By 6 hpi the amplitude of the *α*4 and *α*0 peaks were markedly reduced by both the absence of ICP22 (Fig 5 C and G) and PAA treatment (Fig 5 D and H) when compared to the peaks on the untreated repair virus genome. Because viral DNA replication is inhibited in the ΔICP22 infection at 6 hours, a similar reduction in PPP loading at this time was not unexpected and is consistent with previous results (7).

The potential PPP peak of the *α*22 gene was also reduced in the ΔICP22 infection when compared to the repair infection (Fig 1C). However, this peak contains reads ascribable to PPP of both α22 and *α*47 making its assignment ambiguous. The other *α* gene, *α*27, did not have a well-defined PPP peak in either the repair infection, the ΔICP22 infection, or in wt HSV-1 infection (7), but is similar to the other viral *α* genes with less active Pol II on its locus in the ΔICP22 infection (Fig 4).

### Pol II shuttling between the viral and host genome in the ΔICP22 and repair viruses

We previously demonstrated that from 3 to 6 hpi in wt HSV1-(F) infection, substantial amounts of Pol II are recruited from the host genome to the viral genome (7, 25). To determine the role of ICP22 in this process we first calculated the total, normalized read counts that aligned with either the human or HSV-1 genomes in both infections at each time point (Fig 6 A and B). As expected, the normalized read counts obtained from the repair virus infection aligning with the HSV-1(F) genome significantly increased from 3 to 6 hpi (Fig 6A). This increase occurred concomitantly with a decrease in reads aligning to the human genome in the same infections (Fig 6B).

**Figure 6.**
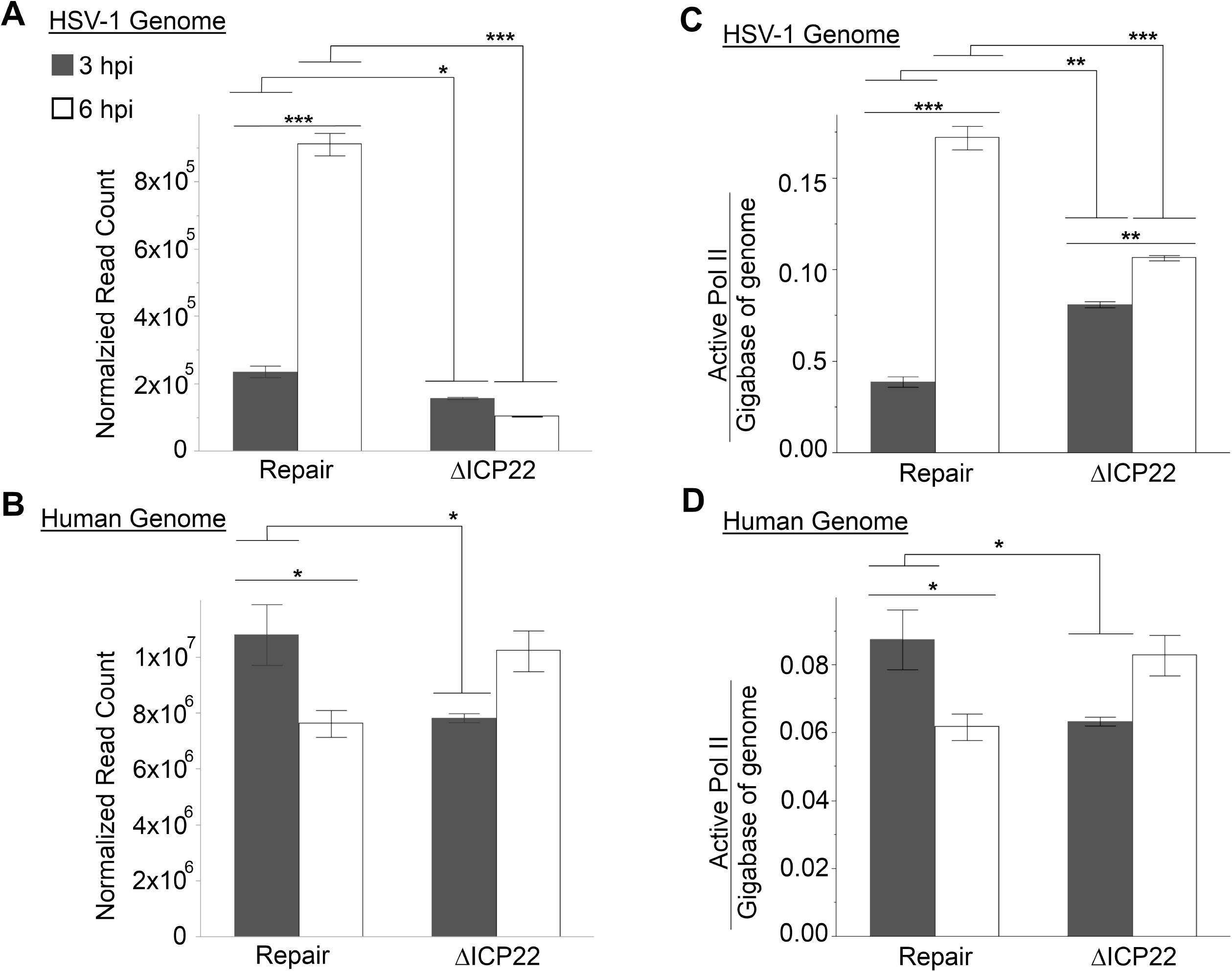
Normalized read count values (**A-B**) and active Pol II molecules per gigabase of DNA template (**C-D**) for the HSV-1 genome (**A and C**) and the human genome (**B and D**) at 3 (gray) and 6 (white) hours post infection with either the Repair or ΔICP22 viruses. The amount of active Pol II/ gigabase of genome value was calculated using the normalized read count divided by the number of estimated nucleotides in question. A two-way ANOVA was used to test for significant differences. A * represents a p-value ≤ 0.05, a ** represents a p-value ≤ 0.01, and a *** represents a p-value ≤ 0.001.

These data indicate that Pol II is redistributed from the human genome to the viral genome over this time period in the repair virus infections, much as we described previously in a PRO-Seq analysis of wild type HSV-1(F).

These results were in stark contrast to those obtained from ΔICP22 infected cells over the same time period. Rather than an increased number of reads aligning to the HSV-1 genome from 3 to 6 hours in the ΔICP22 infection, viral reads actually decreased slightly (p-value = 0.1) over this time (Fig 6A). Moreover, human reads increased over this time period in the ΔICP22 infection, and although this increase was not statistically significant (p-value = 0.067), the result was in stark contrast to the dramatic shift of Poll II to viral genomes in the repair virus infection (Fig 6B). These data suggest that without ICP22, active Pol II does not remain associated with viral DNA, and over time returns to the human genome. To take the effect of viral DNA replication on these results into account, we next calculated the number of active Pol II molecules / bp of DNA template. We divided the number of reads matching a template to the total nucleotide content of that template.

To calculate the amount of template, the 2 × 10^7^ human cells used in each replicate was multiplied by the size of the human genome or the average number of HSV-1 base pairs present at each time point (Fig 3).

As expected, from 3 to 6 hours the amount of active Pol II significantly increased on the repair virus genome (Fig 6A) with a concomitant decrease in normalized reads aligning to the human genome (Fig 6B). The ΔICP22 infection produced 3.16 times less viral DNA template than the repair virus infection (Fig 3), resulting in a greater density of active Pol II per bp of HSV-1 genome in the ΔICP22 infection at 3 hpi than in the repair virus infection (Fig 6C). These data suggest that some of the Pol II molecules occupying the viral genome in the repair infection at 3hpi were located on newly replicated viral DNA.

The density of active Pol II continued to increase from 3 to 6 hpi on the viral DNA in both infections (Fig 6C). However, Pol II occupied the repair virus genome at the greatest density at 6hpi (Fig 6C), and this density was significantly greater than was found on the ΔICP22 viral genome. Thus, without *α*22 and efficient viral DNA replication, Pol II was retained on viral DNA templates at lower densities at later times pi.

### Changes in Pol II activity on cellular genes from 3 to 6 hpi during ΔICP22 or Repair virus infections

Infection with ΔICP22 caused a lower Pol II density on the human genome at 3hpi compared to the repair virus(Fig 6D). Overall, the changes in occupancy over time resulted in similar densities of active Pol II on the human genome at 3hpi in the ΔICP22 as that at 6hpi in the repair infection (Fig 6D). Taken together, these data suggest that ICP22 regulates Pol II activity on the host genome and helps control the shuttling of Pol II from the human genome to the viral genome at early time points post infection.

To further investigate the role that ICP22 plays in host gene transcription during HSV-1 infection, we compared normalized PRO-seq reads from cells infected with both viruses mapping from the TSS to the TTS on 5484 human genes at both time points (Fig 7). Similar to what occurs in wt infection (25), 75% of the host genes in cells infected with the repair virus experienced an overall decrease in normalized read count from 3 hours to 6 hours (Δ in read count < 0) (Fig 7A). These data correlated with the less detailed analysis of host gene transcription presented in Figure 6.

**Figure 7.**
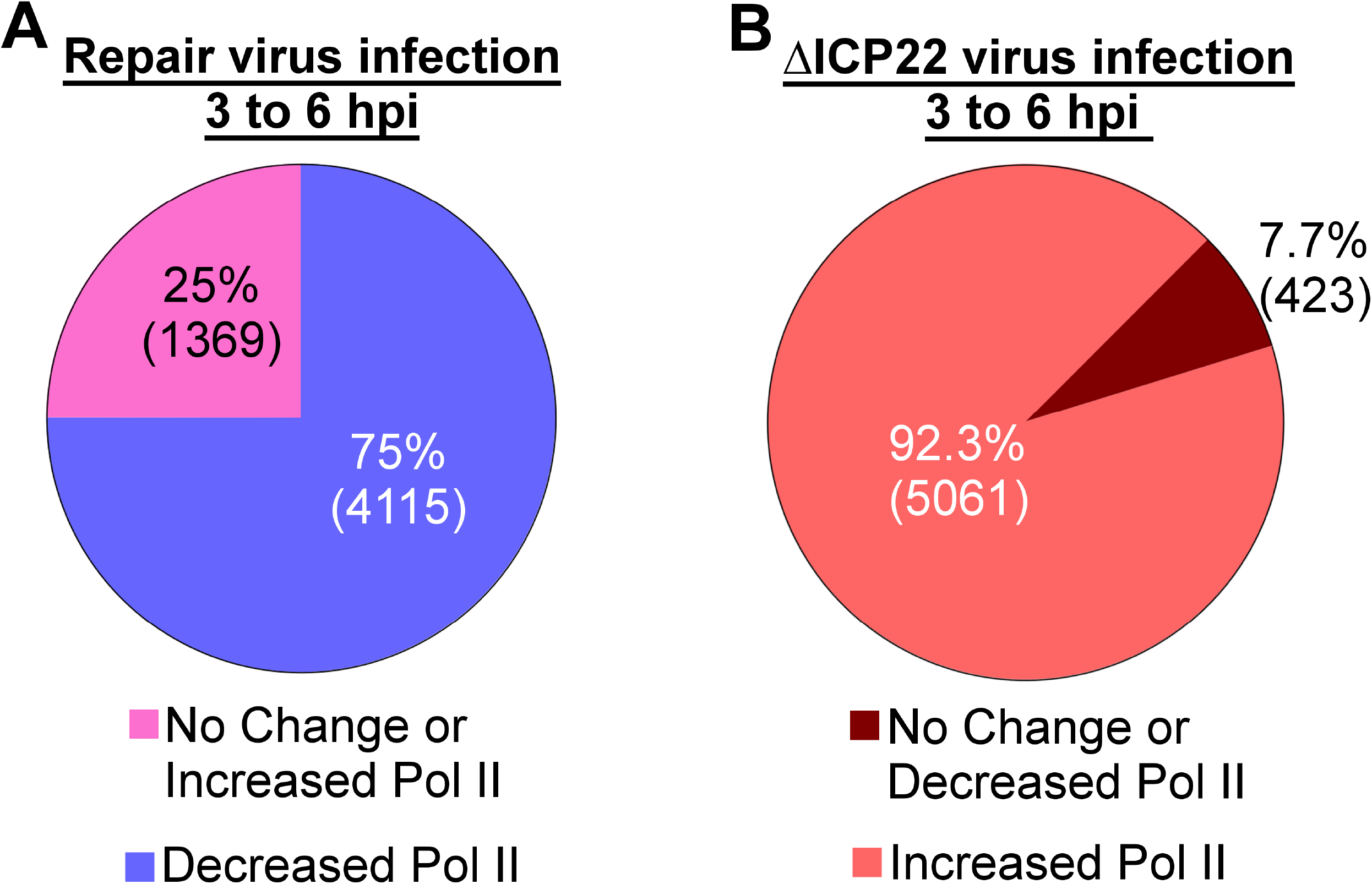
Analysis of the Repair and ΔICP22 infections on human gene transcription. **(A)** The percentage of 5484 transcriptionally active host genes in the repair virus infection that experienced a loss of active Pol II from 3 to 6 hours post infection (Δ in normalized read count < 0) is shown in purple, and those that maintained or increased Pol II activity (Δ in normalized read count ≥ 0) is shown in magenta. (**B)** Changes in Pol II activity on the same 5484 host genes in the ΔICP22 virus infection from 3 to 6 hpi. Coral represents genes that experienced an increase in Pol II (Δ in normalized read count > 0) and dark red represents genes with decreased Pol II activity (Δ in normalized read count ≤ 0).

The same analysis of ΔICP22 infected cells (Fig 7B) produced the opposite phenomenon, with 5061 (92.3%) of the host genes experiencing an increase (a Δ in normalized read count ≥ 0) in active Pol II over the same time period. These data suggest that in the ΔICP22 infection, the majority of the Pol II recruited to the viral genome between 0 and 3 hpi is unable to remain there, and instead migrates back to the host genome in a largely non-specific fashion.

### Log_2_ fold change analysis of active Pol II levels on host genes between the ΔICP22 and Repair virus infections at 3 and 6 hours

When the ΔICP22 induced log_2_ fold change (Supplementary Table 1) for each gene was plotted (Fig 8A) for both the 3-hour time (green) and 6-hour time points (purple), loading on many of the human genes fell between -0.995 and 0.995 (black lines on the y-axis) at both time points. This indicated that a lack of ICP22 during HSV-1 infection does not cause a change in overall Pol II occupancy that is more than 2-fold on most human genes at either time point. However, the ICP22-induced change on most host genes is lower at 3 hours than was seen at 6 hours (as noted by the bolded smooth curves fitting to the data Fig 8A), supporting the conclusion made from Fig 7B, that the return of Pol II to the host genome during the 3-to-6-hour time frame is largely non-specific.

**Table 1.** DAVID database Gene ontology and functional annotation clustering of cellular genes that require ICP22 to maintain active Pol II at their loci in HSV-infected cells at 3 hpi.

| Functional Annotation Cluster | Enrichment Score | Keywords in Classification | Number of Genes | p-value |
| --- | --- | --- | --- | --- |
| **Annotation Cluster 1** | 11.73 |  |  |  |
|  |  | Nucleosome Core | 40 | 7.9E-20 |
|  |  | Chromosome | 47 | 3.9E-7 |
|  |  | Citrullination | 27 | 1.0E-13 |
|  |  | Methylation | 82 | 2.4E-7 |
| **Annotation Cluster 2** | 9.03 |  |  |  |
|  |  | DNA-Binding | 164 | 2.5E-11 |
|  |  | Transcription Regulation | 156 | 1.7E-5 |
|  |  | Transcription | 159 | 2.0E-5 |
|  |  | Nucleus | 328 | 1.2E-11 |
| **Annotation Cluster 3** | 4.87 |  |  |  |
|  |  | Lipid Metabolism | 39 | 7.9E-5 |
|  |  | Cholesterol Metabolism | 13 | 7.3E-6 |
|  |  | Steroid Biosynthesis | 11 | 2.2E-6 |
|  |  | Lipid Biosynthesis | 22 | 7.6E-6 |
|  |  | Sterol Metabolism | 14 | 7.9E-6 |
|  |  | Sterol Biosynthesis | 9 | 9.2E-6 |
|  |  | Cholesterol Biosynthesis | 8 | 1.25E-5 |
|  |  | Steroid Metabolism | 14 | 1.2E-4 |

**Figure 8.**
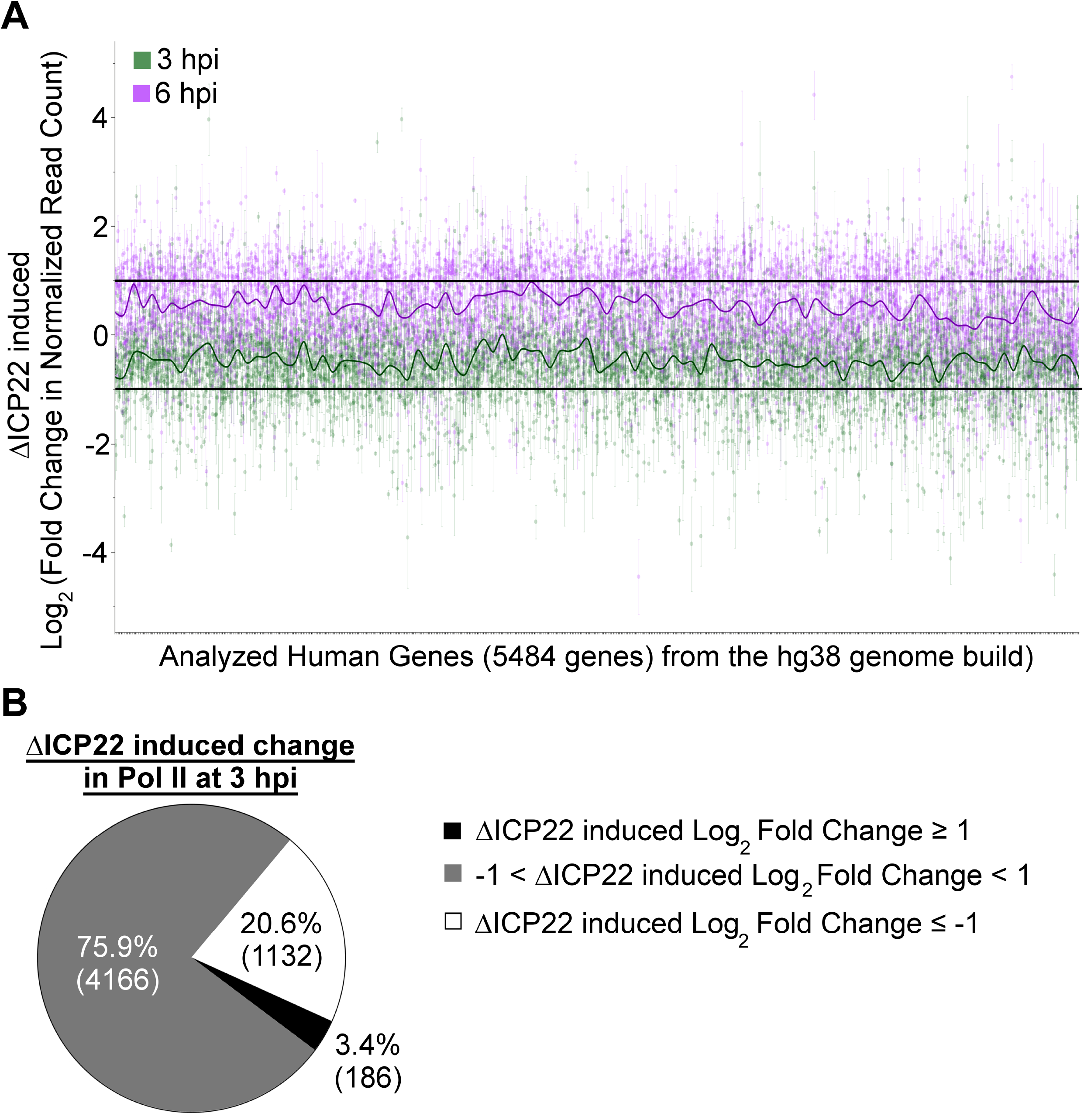
Collinear representation of the ΔICP22 induced log2 fold changes in Pol II activity on the 5484 transcriptionally active host genes at 3hpi (green dots) and at 6hpi (purple dots) aligned with the human genome (**A**). The bolded solid line through the data represents the overall trend in the data. The black lines drawn at 1 and (-) 1 represent cut-offs used to identify the genes shown in (B) for the 3 hour dataset. **(B)** Percentages of host genes at 3 hours pi that either have increased Pol II in the ΔICP22 infection when compared to the repair (log2 fold change in normalized reads ≥0.995) (black), no change in Pol II activity in the ΔICP22 infection when compared to the repair infection (−0.995 < log2 fold change in normalized reads > 0.995) (gray), or decreased Pol II activity in the ΔICP22 infection when compared to the repair infection (a Δ log2 fold change in normalized reads ≤ -0.995) (white).

We were interested in analyzing the 3-hour time point with respect to how Pol II distribution on the human genome was affected by the lack of ICP22. To do this we plotted the % of genes whose ΔICP22-induced log2 fold change that fell in one of three categories defined by the sections of the plot in Figure 8A in a pie chart shown in Figure 8B.

The three categories were defined by the black line cut-offs on the y-axis of Figure 8A, and were: [i] genes with reduced Pol II in the ΔICP22 infection when compared to the repair infection (ΔICP22 induced log_2_ fold change ≤ -0.995), [ii] genes with un-changed Pol II occupancy in the ΔICP22 infection when compared to the repair (−0.995 < ΔICP22 induced log_2_ fold change > 0.995), or [iii] genes with increased Pol II in the ΔICP22 infection when compared to the repair infection (ΔICP22 induced log_2_ fold change ≥ 0.995).

We found that the large majority of host genes at 3 hpi (∼76%) are non-responsive to ICP22, as the ΔICP22 infection produced a less than 2-fold change in read count when compared to the repair virus infection at this time. Very few host genes (186 of the 5484 genes analyzed) had an increase in Pol II in the ΔICP22 infection when compared to the repair virus infection. This suggests that at 3hpi the large shift back to the host genome is either just starting in the ΔICP22 infection or that during the repair infection, ICP22 is acting on a very small subset of genes to promote removal from or addition to the locus.

### ICP22 promotes retention of active Pol II on a subset of host genes

To us, the most interesting genes highlighted by this analysis was the group with less Pol II in the ΔICP22 infection than in the repair (an induced log_2_ fold change in normalized read count ≤ -0.995), indicating a role for ICP22 in Pol II recruitment or retention. This group comprised 1132 host genes (Fig 8B), representing 20.6% of the total human genes analyzed.

To determine if the 1132 human genes that depend on ICP22 for full Pol II occupancy during the first 3 hours of infection shared any functional commonalities, we analyzed the genes with the DAVID Bioinformatics Resources 6.8 (28, 29) supported by the NIH/NIAID. Unexpectedly, this group included a greater than 2-fold reduction in Pol II in 40 of 65 annotated human histone genes (referred to as the nucleosome core in the Up-Keywords database used in the DAVID analysis). These genes were clustered in the first annotation cluster from the DAVID analysis with a fold-enrichment score of 11.73, and had a very significant p-value (Table 1). This analysis indicates Pol II occupancy increases on a subset of human histone genes in a manner dependent on ICP22. Comparing current and previous datasets, we observed that many of the same histone genes had a greater than 2-fold increase in Pol II occupancy after wt HSV-1 infection (25).

Other subsets of genes that showed significant enrichment in this data set are those encoding products involved in transcriptional regulation (specifically the FOXO signaling pathway) and lipid metabolism. As noted previously, many human genes involved in transcriptional regulation maintained Pol II occupancy during wt HSV-1 infection at this time (25). Specific examples that maintained Pol II during wt infection, but lost Pol II after ICP22 deletion include *FOS* and *JUNB* (Supplementary Table 1) (25). These data suggest that ICP22, directly or indirectly maintains Pol II on select host genes early during wt HSV-1 infection.

## Discussion

The mechanism by which ICP22 ensures high levels of active Pol II on α genes is unclear. Previous ChIP experiments using a truncation mutant that likely expresses the first 199 codons of α22 showed ample loading of Pol II on the 5’ ends of viral genes, with relatively less on downstream gene bodies suggesting a role in release to elongation (11). This n199 mutant is able to substantially replicate viral DNA indicating a significant phenotypic difference from the full α22 null mutant (ΔICP22) used in the current study (11). While it is therefore unclear whether the two studies are fully comparable, both studies use mutants lacking ICP22 domains shown to be important for altered Pol II phosphorylation (11, 14, 30) and note decreased Pol II occupancy on immediate early gene bodies (11). Unlike previous work however, we noted decreased activity at α gene PPP sites. The simplest interpretation of our study is that lower levels of transcriptional initiation on immediate early viral gene bodies leads to lower levels downstream. We also cannot formally exclude the possibility that inactive or backtracking Pol II complexes accumulate at immediate early PPP sites in the absence of ICP22 inasmuch as such complexes would not produce a signal in PRO-Seq.

It may be of interest to compare α22 mutants to identify domains responsible for different phenotypes. For example, comparing the current and published data suggest that the N-terminus of ICP22 is important for viral DNA replication. Previous results showed that this region is also important for the formation of virus induced chaperone enriched (VICE) domains that form in the nuclei of infected cells and presumably act to ensure proper folding of intranuclear proteins (31, 32).

In the current study, defects in Pol II activity at the α0 and α4 PPP sites at 3 hours was not phenocopied by inhibition of DNA replication. By 6hpi, however, Pol II activity on the α0, α4 and α27 genes was substantially reduced in both the absence of ICP22 or upon PAA treatment (7). Thus, while ICP22 promotes occupancy of active Pol II on the viral genome at early times, DNA replication plays an important role at later times.

It was of interest to us that in cells infected with ΔICP22 a substantial proportion of Pol II activity that was recruited to viral genes by 3 hpi had returned to cellular genes by 6 hpi. This suggests that ICP22 helps directly or indirectly to retain active Pol II complexes on viral genes, and/or acts to stall or inactivate Pol II complexes on cellular genes, rendering them undetectable by PRO-Seq. The stalling on cellular genes is consistent with the reported ICP22-dependent decrease in Ser 2 phosphorylation of the Pol II CTD, which becomes prevalent at 3-5 hours post infection and continues for the reminder of the infectious cycle (12).

Remarkably, some cellular genes retain or even increase active Pol II after infection (25), and ICP22 is necessary for active Pol II retention for a subset of these genes (Table 1, Supplementary Table 1). This subset included 40 histone genes, including some whose expression is dependent on an activated cell cycle, and some that are expressed constitutively (33, 34). These data support an interesting hypothesis that one function of ICP22 during HSV-1 infection is to increase stable transcripts of host cell cycle-dependent genes whose products are necessary for viral infection (35), without fully activating the cell cycle itself (36-39). Manipulating the cell cycle machinery may contribute to the virus’s ability to engineer the cell for efficient replication in a variety of cell types and conditions.

## Materials and Methods

### Cells and Viruses

The human epithelial lung cancer (HEp-2) cells used for PRO-seq experiments and the African green monkey kidney fibroblast (CV-1) cells used to propagate the HSV-1 (F)-derived viruses were grown in Dulbecco’s modified Eagle’s medium (DMEM) supplemented with 10% new born calf serum (NBS) and penicillin-streptomycin (Pen-Strep) as was done previously (25). The *Drosophila melanogaster* S2 cells used for the PRO-seq spike-in normalization were propagated in Schneider’s medium supplemented with 10% fetal bovine serum FBS as described (25).

The ΔICP22 and matching repair virus were both generated using an HSV-1(F) bacterial artificial chromosome and were kind gifts from Dr. Yasushi Kawaguchi (21, 24). Stocks of both viruses were grown in CV-1 cells. Because the ΔICP22 virus produced poorly defined plaques on cell monolayers, infectious virus in viral stocks was quantified using a fluorescent plaque assay. Briefly, CV-1 cells were infected with serially diluted virus in 199X medium supplemented with 1% NBS and Pen-Strep for 2 hours at 37°C. At this time, the medium was changed to DMEM supplemented with 2% NBS and Pen-Strep and the infection was allowed to continue at 37°C for 72 hours. Media was removed and cells were fixed with methanol and blocked with 1% bovine serum albumin (BSA) before being incubated for 2 hours at room temperature with a 1° rabbit antibody against HSV-1 glycoprotein M (g(M)) diluted 1:1000 in PBS plus 1% BSA (40). The cells were then washed and incubated for 30 minutes at room temperature with a 2° goat anti-rabbit antibody labeled with alexafluor 488 that was diluted 1:5000 in PBS plus 1% BSA before washing with excess PBS. Plates underwent a final, brief wash with H_2_O before being dried and stored in the dark until plaque counting. Fluorescent plaques were observed and counted using an Olympus IX71 fluorescence microscope to determine infectious viral titer. The repair virus was quantified using a crystal violet-stained CV-1 cells.

### Virus Infections and Drug Treatments

For one PRO-seq replicate, 2 × 10^7^ HEp-2 cells were infected with either the ΔICP22 or repair virus at an MOI of 5 in 199V medium (199X medium supplemented with 1%NBS and Pen-Strep) for 2 hours at 37°C. Media was then changed to DMEM 2% NBS and Pen-Strep and the infection was allowed to continue to either 3 or 6 hpi. For the DNA replication experiment, 2 × 10^6^ HEp-2 cells were infected at an MOI of 5 in the same manner as was done for the PRO-seq experiments. In some experiments, cells infected with the repair virus were treated with PAA. This was done by adding PAA to a final concentration of 0.2 mg/ml to the 199V and DMEM 2% NBS medium used for the infections, similar to what has been described previously (7).

### Nuclei Isolation and Cytoplasmic Extract Preparation

For the PRO-seq analysis, HEp-2 cell nuclei were isolated on ice by exposing the washed, adherent cells to a swelling buffer [10 mM Tris-HCL (pH 7.5)], 10% glycerol, 3 mM CaCl_2_, 2 mM MgCl_2_, 0.5 mM dithiothreitol [DTT], and 4 U/ml RNase inhibitor) before removing them from the plate, and exposing them to a lysis buffer (10 mM Tris-HCL [pH 7.5], 10% glycerol, 0.5% Igepal, 3 mM CaCl_2_, 2 mM MgCl_2_, 0.5 mM DTT, and 4 U/ml RNase inhibitor) as previously described (7, 10, 25, 41). After lysis, the nuclei were collected by centrifugation at 1500 x g for 5 minutes at 4 °C before being washed twice with ice cold lysis buffer and one time with freezing buffer (50 mM Tris-HCL [pH 8], 25% glycerol, 5 mM Mg(CH_3_COO)_2_, 0.1 mM EDTA, 0.5 mM DTT, and 4 U/ml RNase inhibitor) before being resuspended at a final density of 2 × 10^7^ nuclei/ 100 μl of freezing buffer before being flash frozen in liquid N2.

The Drosophila melanogaster S2 cell nuclei were isolated in the same manner but were resuspended at a final density of 2000 nuclei/ 25 ul of freezing buffer before being flash frozen in liquid N2.

### Precision Nuclear Run-On assay

Run-on reaction conditions followed previously published protocols (7, 22, 25). The frozen HEp-2 nuclei and S2 nuclei were defrosted on ice, and S2 nuclei were spiked into the HEp-2 nuclei at a final ratio of 1 S2 nucleus : 1000 HEp-2 nuclei prior to the addition of the run-on reaction buffer (10 mM Tris-HCL [pH 8], 300 mM KCL, 1% Sarkosyl, 5 mM MgCl_2_ 1 mM DTT, 100 um biotin-11 ATP, 100 um biotin-11 CTP, 100 um biotin-11 GTP, 100 um biotin-11 UTP, and 4 U/ml RNase inhibitor). Nuclei were thoroughly mixed with the run-on reaction buffer and the reaction was allowed to proceed for 3 min at 37° C with constant shaking on a vortex shaker with a speed setting of 8. The run-ons were terminated with the addition of 500 ul of TRIzol™ LS followed by RNA extraction.

### Sequencing Library Preparations

The RNA from the precision nuclear run-on was hydrolyzed with 0.2 N NaOH for 20 min before excess nucleotide removal with a BioRad P-30 column. Biotinylated transcripts were purified from the total RNA pool using M280 streptavidin Dynabeads and a series of washes consisting of a high salt (50 mM Tris-HCL [pH 7.4], 2 M NaCl, 0.5% Triton X-100, and 2 U/ 10 ml RNase inhibitor) a medium salt (10 mM Tris-HCL [pH 7.4], 300 mM NaCl, 0.1% Triton X-100, and 2 U/ 10 ml RNase inhibitor), and low salt (5 mM Tris-HCL [pH 7.4], 0.1% Triton X-100, and 2 U/ 10 mL RNase inhibitor) before being extracted in Trizol™ as described previously (7, 25).

Once purified, the 3’ end of the biotinylated RNA’s were ligated to a 3’ RNA adapter containing a 5’ phosphate group (5’-Phos) and a 3’ inverted dT (InvdT) (5’-Phos-GAUCGUCGGACUGUAGAACUCUGAAC-3’-InvdT) with T4 RNA ligase I. After a subsequent round of bead binding and washing, the 5’ 7-methylguanosine caps of the biotinylated RNAs were removed with 10 U of 5’-pyrophosphohydrolase (RppH) and the 5’ ends repaired using T4 polynucleotide kinase. The 5’ end of the RNAs were then ligated to the 5’ RNA adapter (5’-CCUUGGCACCCGAGAAUUCCA-3’) with T4 RNA ligase I, and subjected to a third round of bead binding, washing, and extraction.

The resulting RNA was reverse transcribed using SuperScript III reverse transcriptase (Thermo Fisher), the RNA PCR primer 1 (5’-AATGATACGGCGACCACCGAGATCTACACGTTCAGAGTTCTACAGTCCGA-3’), and a 0.625mM concentration of each of the four deoxyribonucleotides, 5mM DTT, and 20 U of RNase inhibitor in the provided buffer. Two microliters of cDNA were removed from each sample and used for a test PCR amplification to determine the optimum number of PCR amplification cycles needed for each sample set. Once this value was determined the remaining cDNA was PCR amplified for the determined number of cycles using the Phusion high-fidelity DNA polymerase and the provided 1x GC buffer from New England Biolabs, and 0.25 *μ*M of the barcoded Illumina RNA PCR index primer and 0.25 mM concentration of each of the four deoxyribonucleotides. The resulting sequencing libraries were purified on an 8% polyacrylamide gel in 0.5 x Tris-borate-EDTA (TBE) and the library fragments above 120 nucleotides in length were extracted from the gel and submitted for sequencing on the Illumina NextSeq 500 platform at the GeneLab sequencing facility at the Louisiana State University School of Veterinary Medicine.

### Bioinformatics and Statistical Analysis

The raw sequence reads were processed with the PRO-seq pipeline developed by Charles Danko and colleagues (https://github.com/Danko-Lab) as previously described (7, 25). This process cuts off the adapter sequences from the raw reads and aligns them to a concatenated genome containing the human hg38 genome build, the drosophila dm3 genome build, the HSV-1 F genome build, and the rRNA genome build which reduces misalignment of similar reads from different genomes. The output .BAM files were then used to probe the drosophila genome for normalization, or for individual viral and human genes and gene regions with the SeqMonk software produced and maintained by the Babraham Institute (https://www.bioinformatics.babraham.ac.uk/projects/seqmonk/) (27).

The Drosophila normalization read count values were obtained using a running-window analysis of total reads from both strands of the drosophila genome, which gave the values used for the normalization. The human and viral genes were probed for reads occupying the whole gene (TSS to TTS). The viral reads were also probed for reads falling in the PPP region (TSS to +100 nt downstream), or mapping in the first 100nt body region (+100nt downstream of the TSS to +200 nt downstream of the TSS). The output .bw files were used to view Pol II occupancy over an entire gene or gene region with the integrative genomics viewer (IGV) from the Broad Institute (http://software.broadinstitute.org/software/igv/) (26, 42), using the drosophila normalization values to appropriately adjust the viewing scale of the data.

The .bw files were also used in conjunction with the pyBigWig program to extract the raw read data for each nucleotide in the first 200 nt region of the *α*4 and *α*0 gene from each sample. The reads were normalized with the normalization factor derived from the drosophila spike in and plotted using the smoother function of the graph builder in the SAS JMP Pro software for Macintosh on x64, JMP^®^, Version 15.1.0. SAS Institute Inc., Cary, NC, 1989-2019. The confidence of fit was calculated using bootstrapping with 300 different iterations of the data.

The drosophila-normalized read counts for the probed regions were analyzed using the SAS JMP Pro software for Macintosh on x64, JMP^®^, Version 15.1.0. SAS Institute Inc., Cary, NC, 1989-2019 using the biological replicates and analyses indicated for each figure. In brief, depending on the questions being addressed, the data was compared using a one-way analysis of variance (ANOVA), two-way ANOVA, or 3 way-factorial analyses with a student T-test or Tukey honestly significant difference (Tukey’s HSD) post-test with an ordered-difference report to produce p-values for the comparisons being made.

### Host gene analysis

The BAM files from the sequence alignment to the hg38 genome build were used for analysis of Pol II activity on the host genome. The raw reads were subjected to a DESEq2 analysis using SeqMonk to identify host genes that were significantly represented in the data set. The raw reads from these identified genes were then normalized using the drosophila spike-in and used for the log2 fold change calculations.

The functional annotation clustering analysis was performed using the DAVID Bioinformatics Resources 6.8 (28, 29) software supported by the NIH/NIAID. The up-keywords database was used in the clustering with a medium classification stringency.

### Viral DNA Replication Assay

The amount of viral DNA replication in the repair and ΔICP22 virus infections was analyzed using absolute quantitative PCR analysis of the number of viral genomes existing in 50 ng of DNA extracted from the described, infected HeP-2 cells as done previously (7, 25, 43). DNA was extracted from HeP-2 cells at the indicated time using the GeneJET genomic DNA purification kit from Thermo Fisher^®^. The DNA concentration was determined by spectrophotometry, and 50 ng was analyzed in a Sybr-green based qPCR assay using primers specific to the HSV-1 UL51 gene and the Bio-Rad Sybr Green master mix in conjunction with a standard curve of serially diluted pcDNA3 plasmid containing an insert of the UL51 gene.

## Data Availability

All sequencing data including the raw FASTq files from the Illumina Next Seq 500 sequencing run, and the processed bigwig files are available on the GEO database. The data is currently private but will be made publicly available upon publication. Currently the data can be viewed by peer reviewers at the GEO database website (https://www.ncbi.nlm.nih.gov/geo/) with the accession number: GSE169574, and the peer reviewer secure token: ypwzksmszpcrtej

## Acknowledgments

We thank Dr. Yasushi Kawaguchi for the ICP22 deletion and repair viruses, Thaya Soufflet and Dr. Valdimir Chouljenko at the BIOMMED division at the LSU School of

Veterinary Medicine sequencing facility for their help with the NextSeq500, and Dr. Xue Wen at the LSU School of Veterinary Medicine for help with the statistical analysis. The high-throughput sequencing portion of this research was performed using the SuperMic supercomputer provided and maintained by the Louisiana State University high performance computing department (https://www.hpc.lsu.edu). Specifically, we thank Dr. Le Yan for his help with maintaining the PRO-seq sequencing pipeline and for installing the pyBigWig program. JDB is a Scientific Advisory Board Member for Virios, which neither reviewed nor funded the current study. These studies were supported by National Institutes of Health grants R01 AI 141968 and R21 AI 148926 to JDB.

## Figure Legends

**Table 1**. Significant results from the DAVID Gene ontology functional annotation clustering of host genes that require ICP22 to maintain active Pol II at their loci at 3 hpi. The first three functional annotations are shown along with their enrichment scores. The keywords that fit into each classification are indicated below each cluster, along with the number of genes found in the dataset that match the keyword and the corresponding p-value representing the significance of this enrichment within the datset.

**Supplementary Table 1**. Log2 fold change calculations and standard deviations of the ΔICP22 induced changes in Pol II activity at 3 and 6 hpi on the 5484 host genes found to be significantly enriched in the data set using a DESEQ2 analysis. Calculations were made by averaging the normalized read counts of two independent biological replicates after the DESEQ2 analysis for significant enrichment in the dataset. The human genes are referred to both by their name and their Ensemble Gene ID.

